# Mineral analysis of complete dog and cat foods in the UK and compliance with European guidelines

**DOI:** 10.1101/172544

**Authors:** Davies M., Jones L., Alborough R., Davis C., Williams C., Gardner D.S.

## Abstract

The mineral content of complete pet food is regulated to ensure health of the companion animal population. A comprehensive analysis of adherence to these regulatory guidelines has not been conducted. We measured mineral composition of a range of complete wet (n=97) and dry (n=80) canine and feline pet food sold in the UK to assess compliance with EU guidelines. While a majority of foods complied with ≥8 of 11 guidelines (99% and 83% for dry and wet food, respectively), many failed to provide nutritional minimum (e.g. Cu, 20 % of wet food) or exceeded nutritional maximum (e.g. Se, 76% of wet food). Only 6% (6/97) of wet and 39% (34/80) of dry food were fully compliant. Some foods (20-30% of all analysed) had mineral imbalances such as not having the recommended balance of Ca:P (between 1:1 to 2:1). Foods with high fish content had high levels of undesirable metal elements such as arsenic. The study highlights broad non-compliance of a range of popular pet foods sold in the UK with EU guidelines (95% and 61% of wet and dry foods, respectively). If fed exclusively and over an extended period, a number of these pet foods could impact the general health of companion animals.

## Introduction

Companion animals, particularly dogs and cats, have requirements for essential minerals that need to be supplied in their food (NRC, 2006). In the UK, wet pet foods (e.g. tins, pouches) are fed to 41% of dogs and 77% of cats (PFMA, 2016), usually together with treats, snacks (including table scraps) or other complementary foods. Thus, more than half of the companion animal population only consume the main ration provided by their owners, with the remainder consuming a mixed, likely nutritionally-unbalanced diet. In a pan-European Harris poll, 56% of UK pet owners bought branded pet foods, higher than any other market in Europe (Grocer, 2010). Brand loyalty often results in the same product being fed exclusively to the pet for long periods of time (months to years). It is therefore imperative that such branded wet and dry foods, labelled a ‘complete’ are nutritionally replete and balanced for macro- and micronutrients including sufficient, but not excessive, essential mineral elements.

In countries of the European Union, guidelines established by the Fédération Européenne De I’industrie des aliments pour Animaux Familiers (FEDIAF, the European Pet Food Industry Federation) provides pet food companies with recommended concentrations of macronutrients (e.g. crude protein, fat), micronutrients (e.g. specific vitamins and minerals) and amino acids (e.g. arginine, lysine) (FEDIAF, 2013). Levels of incorporation or ‘daily allowance’ within a diet for each nutrient are defined as ‘*the level of intake of a nutrient or food component that appears to be adequate to meet the known nutritional needs of practically all healthy individuals. It reflects the minimum requirement plus a safety margin for differences in availability between individual animals and for nutrient interactions. In practice this would be translated as the levels of essential nutrients that healthy individuals should consume over time to ensure adequate and safe nutrition*’. Such foods should therefore have the correct proportions of essential macro- and micronutrients that are sufficient for a daily ration (EU Regulation No 767/2009; art. 3(i), adapted to pet food) with a daily ration satisfying all of an animal’s energy and nutrient requirements without further complementary intake. These EU guidelines were largely based upon the original National Research Council recommendations in the USA, but for many minerals there is no specific recommendation and no ‘safe upper limit (SUL)’, i.e. above which the micronutrients can become toxic (NRC, 2006). For toxic elements such as arsenic, cadmium, mercury and lead, further legislation regarding acceptable concentrations in foods are regulated by EU directive 2002/32/EC.

To our knowledge only one previous study, conducted in the USA, assessed total mineral content of 45 dry foods for dogs with reference to American recommendations (AAFCO; (Gagne et al., 2013)). Thirteen percent did not comply with AAFCO guidelines for certain minerals and many diets had excessive calcium. Long-term feeding of un-balanced or of diets deficient or excessive in certain minerals can have adverse impacts on animal health (Davies, 2016, McDowell, 1992). For this reason, we assessed, for the first time in the European Union, mineral composition of a broad range of dry and wet foods for consumption by domestic dogs or cats. We referenced composition against 11 o 13 current EU guidelines for minerals (FEDIAF 2013). Briefly, we find widespread non-compliance; only 6% and 39% of wet and dry foods, respectively complied with 11 of 13 FEDIAF guidelines. Many individual products were mineral imbalanced (e.g. Ca:P ratio as low as 0.3:1 or as high as 3:1) or had high concentrations of individual minerals (e.g. selenium ≥300 μg 100g DM^-1^; arsenic ≥1.1 mg 100g DM^-1^) that if fed exclusively for many months could impact the general health of companion animals.

## Materials and Methods

### Ethics

This study was approved by the University of Nottingham School of Veterinary Medicine and Science Research Ethics Committee.

### Selection of Pet foods

Pet foods were bought locally from a range of commercial suppliers and pet food supermarkets and were packaged in a mixture of sealed tins, pouches, cans or sealed bags. A total of 177 different pet foods were included for analysis, all labelled as ‘Complete’. The foods were selected as representative of popular brands sold in the UK and including a range of main flavours (e.g. beef, chicken, fish) largely aimed at adult pets (139 of 177) but also young (n=14) or senior (n=24) animals. In total, 42 different brands were represented (dry food, n=21 brands; wet food, n=27 brands) with 113 products for cats (wet food, n=48; dry food, n=65) and 64 products for dogs (wet food, n=49; dry food, n=15). Key nutritional information provided on the label was recorded such as macronutrient content (percentage protein, fat, moisture, ash and fibre, as fed) alongside the country of origin and batch number. Each product was designated as either a ‘supermarket own-brand’ food or a ‘prescription/veterinary/therapeutic’ diet based upon the label and target population. Other than these categories, all other products were designated as a ‘standard’ feline or canine food. Energy content (gross, digestible, metabolic) was rarely provided on the label and was therefore calculated using modified Atwater criteria (Equations 1-7; Supplementary Information). The concentration of minerals added to each diet was also recorded, if provided (e.g. additional zinc, copper and iron; all as μg/kg). Further information on the composition of the food was derived according to the predominant labelling and checked according to the order of primary ingredients, e.g. main protein source being beef, chicken or fish.

### Assessment of compliance

Complete foods should have, as defined by the EU, the correct proportions of essential macro- and micronutrients that is sufficient for a daily ration (EU Regulation No 767/2009; art. 3(i), adapted to pet food). The daily ration should satisfy all of an animal’s energy and nutrient requirements without further complementary intake (FEDIAF guidelines for dogs in Table S1). These guidelines were largely based upon the original National Research Council minimum recommendations in the USA (Table S2). Nevertheless, for many minerals there is no specific recommendation and no ‘safe upper limit (SUL)’, i.e. above which the micronutrients can become toxic (NRC, 2006). In the United States of America, pet food manufacturers also receive guidance from the American Association of Feed Control Officials for maximum concentrations in pet food (Table S3; (AAFCO, 2015)). In the EU, aside from guideline quantities of macronutrients (crude protein, fat, ash etc…) there are thirteen specific recommendations for macrominerals (calcium, phosphate, Ca:P, potassium, sodium, chloride and magnesium) and trace minerals (copper, iodine, iron, manganese, selenium and zinc). In addition, recommended maximum levels of ‘undesirable’ or heavy elements (arsenic, cadmium, mercury and lead) in pet food are regulated by EU directive 2002/32/EC (Table S4.

### Preparation of samples for elemental analysis

To avoid trace ion contamination of the samples, plastic utensils were used when preparing all food stuff and gloves were worn at all times. A known, representative quantity (100-200g) of wet food (for small pouches and tins, all contents) or dry food was first opened then emptied into a 250ml solvent-resistant container (Sarstedt, UK). Contents were then freeze dried for 2 days and re-weighed to establish moisture content by difference. For larger cans and dishes of wet food (≥500g), the whole sample was pre-mixed and a representative sample (100-200g) was obtained for freeze-drying. Dried foods were ground to a consistent powder and duplicate samples (100-200 mg) of this powder were acid-digested using standard techniques for mineral analysis by inductively-coupled plasma mass-spectrometry (ICP-MS), a method widely considered the gold-standard for mineral analyses of various fluids, tissues and other biological composites such as foodstuffs (Goullé et al., 2005, Joy et al., 2015). In brief, after 1 hour incubation in 3.0ml of 68% trace analysis grade (TAG) HN0_3_, 2.0ml 30% H_2_O_2_ (both Fisher Scientific UK Ltd, Loughborough, UK), and 3.0ml milli-Q water (18.2 MΩ cm), acid-digestion was accelerated by microwave-heating (Anton-Paar, 2000) for 45mins in 3.0 ml of 70% Trace Analysis Grade (TAG) HNO3, 2.0 ml H_2_O_2_ and 3.0 ml milli-Q water (18.2 MΩ cm) (Fisher Scientific UK Ltd, Loughborough, UK). The resulting solution was diluted to 15ml with milli-Q water. To control for unexpected sources of contamination (e.g. in acids or water) duplicate blank tubes were run with each batch (containing all liquids but no sample). Additionally, for every batch (to control for batch-to-batch variability) and/or every 60 ICP-MS tubes (control for within-run drift), duplicate samples of a certified reference material (CRM) were included. For all food digests, the CRM was NIST (National Institute of Standards and Technology, Gaithersburg, MD, USA) 1577c (bovine liver). All single batch data for individual elements were corrected to percentage recovery of the CRM (Table S5). Minimal between-batch variability was accounted for in statistical models by including BatchID as a random effect. Data for minerals not included on 1577c and thus not subjected to internal quality control were measured, but are not reported (B, Li, Be, Al, Ba, TI and U). Percentage recovery for nickel was relatively low and is also not reported.

### Elemental analysis by ICP-MS

Elemental analysis was by inductively coupled plasma-mass spectrometry (ICP-MS; iCAPTM Q, Thermo Fisher Scientific Inc., Waltham, MA, USA) using a He collision cell with ‘kinetic energy discrimination’ to reduce polyatomic interference in the analysis of Ag, Al, As, B, Ba, Cd, Ca, Co, Cr, Cs, Cu, Fe, K, Mg, Mn, Mo, Na, Ni, Pb, Rb, S, Sr, Tl, U, V and Zn. Lithium, Be and P were determined in standard (vacuum) mode and Se in ‘hydrogen-cell’ mode, with ‘insample switching’. Internal standards were Ge, Rh and Ir. Final foodstuff elemental composition is presented after correction for blanks and batch variation (using the CRM as reference) as ppm (e.g. major elements, 1ppm = 1mg/kg), ppb (e.g. trace elements, 1ppb = 1μg/kg) or per 100g of dry matter [DM] as indicated in the text. For major and trace elements, recovery was >95% with <10% coefficient of variation for each (n=17 separate analyses). Intra-assay variability for all elements was <2%. Beryllium (Be), Lithium (Li), Silver (Ag), Titanium (Ti), Thallium (TI) and Uranium (U) were identifiable but the majority of results were close to limits of detection (LOD) and are not reported. Lead (Pb), Caesium (Cs), Cadmium (Cd) were above LOD but below limits of quantification (LOQ) and are therefore also not shown. Chromium (Cr), Molybdenum (Mo) and Vanadium (V) were above LOQ but many results were close to LOQ and therefore these data are also not included.

## Results

### Standardisation of mineral detection

Analysis of 12 veterinary diets with a stated guaranteed analysis for 5 minerals were compared against values determined by ICP-MS at The University of Nottingham, as described above. Excellent comparisons were achieved with a highly-significant correlation between estimates of mineral composition (Figure S1). Furthermore, a number of food samples were analysed repeatedly within-run (8 technical replications) yielding a coefficient of variation of <5% for elements at low (e.g. arsenic) or high (e.g. selenium) concentration.

### Macronutrient and energy composition of the diets

As expected, foods fed to cats had greater crude protein content and lower carbohydrate, than foods fed to dogs (Table 1). Total mineral content (reflected as declared ash on the label) also tended to be increased in foods fed to cats (Table 1). On an as-fed basis, fat content was similar between wet foods, but increased in dry foods fed to cats versus dogs (Table 1). Foods fed to cats had higher mean energy density per se, but after correction for dry matter content, were similar (Table 1). The declared ash content of pet food (corrected to DM) correlated well with measured total mineral content, but considerable variability was noted particularly for ‘standard’ wet diets (Figure S2). Foods designated as ‘supermarket own brand’ tended to have higher ash content than ‘prescription or therapeutic’ diets (Figure S2).

**Table 1:**
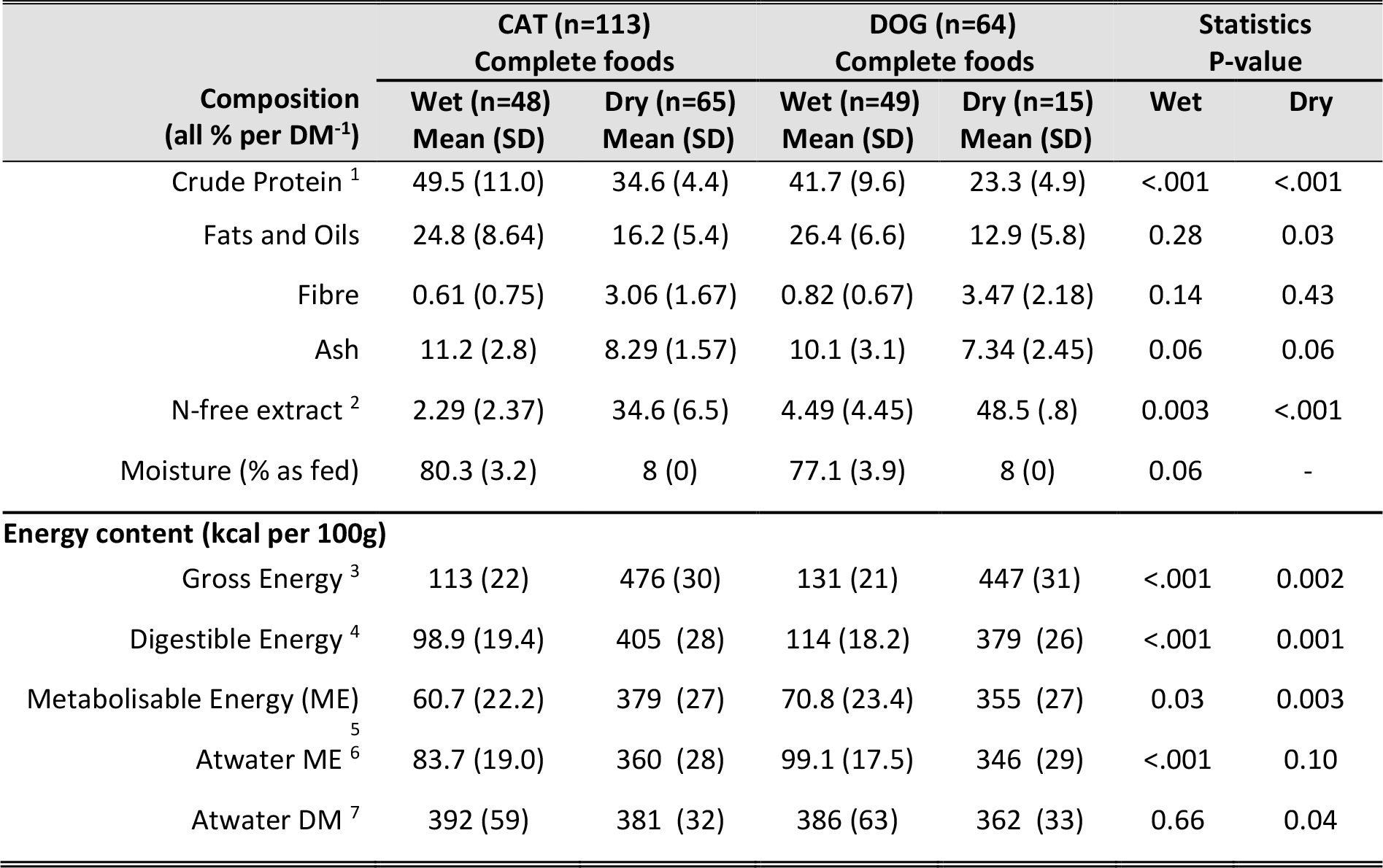
Macronutrient and calculated energy content of complete cat and dog food. Macronutrient content were derived from the stated composition on the label each product. Products were clearly labelled as being a complete diet, for a cat or dog and were either wet food or dry kibble. Energy densities were calculated according to modified Atwater criteria as outlined in Equations 1-7, supplementary information. All data were normally distributed and were analysed by 1-way ANOVA within each type of food to test for differences in composition of foods fed to cats or dogs. A formal comparison between wet or dry food was not considered of value. A P-value of <0.05 was accepted as indicating statistical significance.

### Mineral (elemental) composition of the diets

Overall elemental composition of foods for cats or dogs was broadly similar (i.e. no significant difference for total major or trace elemental composition; Table 2). Corrected for moisture content, wet pet food had higher sulphur, calcium, phosphorus, sodium, potassium, strontium and rubidium but lower manganese relative to dry food (Table 2). Foods (wet or dry) fed to cats had lower magnesium but higher potassium than foods fed to dogs (Table 2). When all foods were represented according to the mandatory FEDIAF guideline (Table S1, 11 of 13 guidelines were tested; chloride and iodine were not measured) some interesting outliers were noted: for example, two foods had Ca:P ≤ 0.25 whereas three others had a Ca:P ratio of ≥ 2.5 (Figure 1a). Most foods complied with guidelines for elemental iron due largely to the exceptionally wide acceptable range (Figure 1b). A number of foods, particularly wet food, were either above or below FEDIAF guidelines for copper (Figure 1c) and the majority of wet foods exceeded the legal maximum for selenium (Table S1; 56.8 μg/100g DM), which had a very narrow acceptable range (Figure 1d).

**Figure 1:**
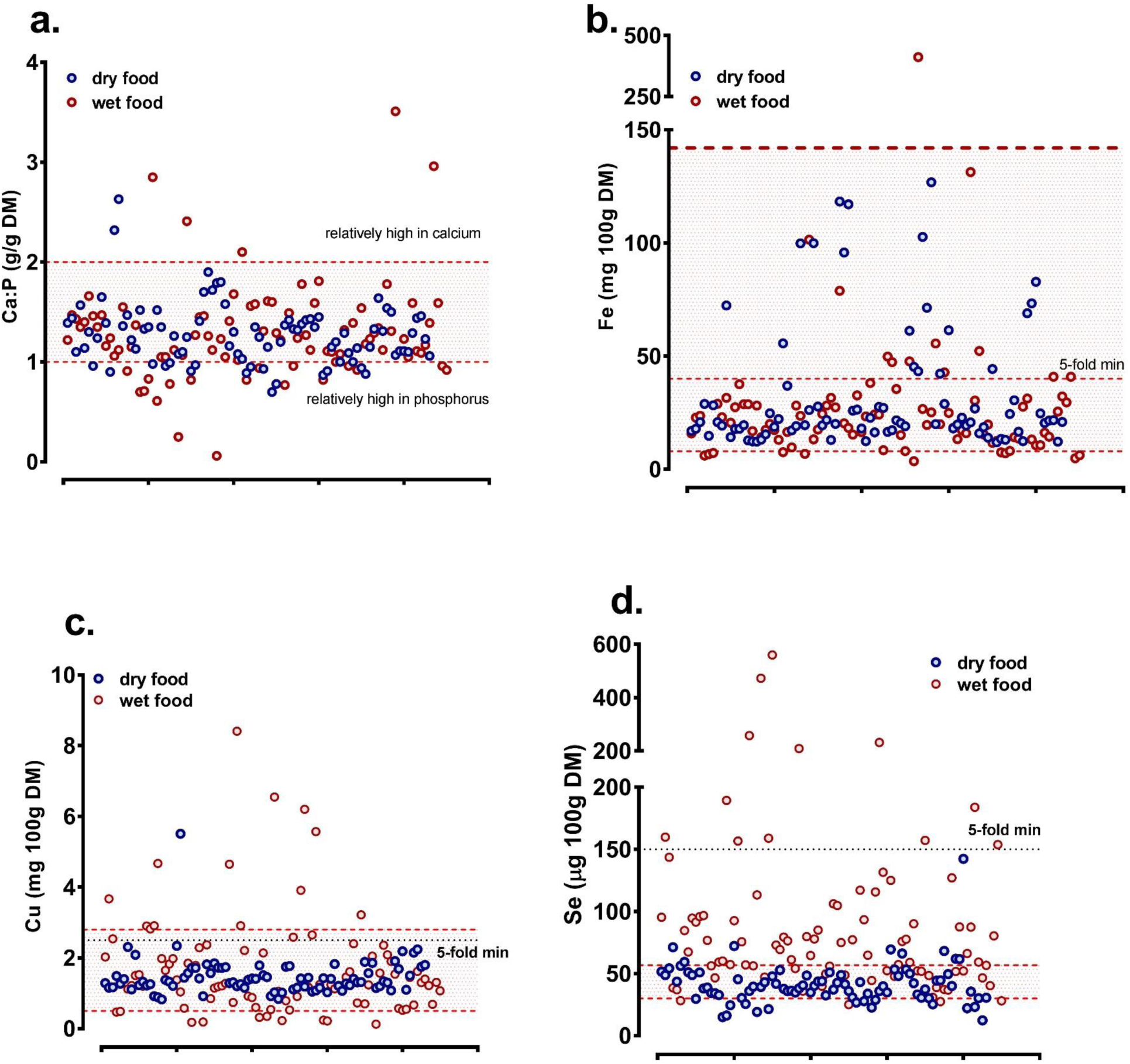
Calcium to Phosphorus, Copper, Iron and Selenium content of complete, wet and dry pet foods. **a-d**. Total elemental matter (Calcium [Ca], Phosphorus [P], Iron, [Fe], Copper, [Cu] and Selenium, [Se]) were measured in duplicate samples of freeze-dried, homogenized and nitric acid-digested pet food by ICP-MS. Each dot represents a single wet (red shaded dots) or dry (blue shaded dots) food. Lower and upper dotted lines represent nutritional minimum and maximum, respectively as recommended by FEDIAF. Black dotted line represents incorporation at 5-fold nutritional minimum, since few recommendations for a feline nutritional maximum exists. Data for each product are plotted individually along the x-axis, with elemental density on the y-axis.

**Table 2:** Elemental composition of complete, wet cat and dog food. Data are presented as median (IQR; 1^st^ to 3^rd^ quartile) for each element. Duplicate, freeze-dried homogenized samples (100-200mgs) were analysed by ICP-MS, and a mean value established. Since elemental composition is presented as parts (ppm, ppb) per unit dry matter, a formal statistical comparison was conducted as a 2 (cat vs dog) × 2 (wet vs dry) factorial ANOVA, with the interaction term included. If necessary, data were logi0 transformed prior to analysis in order to normalize residual errors, to avoid distributional bias. Statistical significance was accepted at P<0.003 (Bonferroni correction for 14 elements above LOQ). Data for total trace elements includes As, Co, Cr, Cs, Cu, Fe, Mn, Mo, Rb, Se, Sr, V and Zn.

| Major element<br>(all ppm, mg/kg DM) | CAT (n=113)<br>Complete foods |  | DOG (n=64)<br>Complete foods |  | Statistics<br>P-value |  |
| --- | --- | --- | --- | --- | --- | --- |
|  | WET (n=48)<br>Median (IQR) | DRY (n=65)<br>Median (IQR) | WET (n=49)<br>Median (IQR) | DRY (n=15)<br>Median (IQR) | Cat vs.<br>Dog | Wet vs.<br>Dry |
| Sulphur | 7269 (6348,8442) | 4989 (4124,6447) | 5211 (4442,7528) | 3302 (2925,3923) | 0.01 | <.001 |
| Calcium | 17423 (13580,25565) | 14125 (11442,19456) | 19766 (15147,27387) | 12860 (6290,17813) | 0.21 | <.001 |
| Phosphorus | 15715 (13297,19300) | 12365 (10730,14194) | 15672 (10471,21473) | 9555 (5051,13016) | 0.59 | <.001 |
| Sodium | 11906 (6437,17135) | 5692 (4233,6319) | 10180 (6350,14148) | 4093 (2743,4375) | 0.63 | <.001 |
| Magnesium | 1048 (847,1195) | 1110 (915,1357) | 1353 (1080,1762) | 1208 (1129,1360) | <.001 | 0.86 |
| Potassium | 13376 (9943,18670) | 7686 (7174,8615) | 14857 (10102,19474) | 6254 (5567,7593) | <.001 | <.001 |
| **TOTAL MAJOR** | 73070 (61513,82443) | 47352 (41733,54338) | 70925 (54602,86870) | 41224 (25416,46955) | 0.07 | <.001 |
| **Trace element (all ppb, µg/kg DM)** |  |  |  |  |  |  |
| Zinc | 154 (127,178) | 176 (143,208) | 170 (137,223) | 180 (145,214) | 0.03 | 0.21 |
| Manganese | 19.7 (11.3,24.3) | 59.6 (37.8,69.4) | 23.3 (15.9,35.4) | 57.9 (36.9,75.7) | 0.05 | <.001 |
| Strontium | 15.0 (10.2,33.8) | 13.9 (11.4,17.0) | 20.2 (14.0,27.6) | 10.9 (9.34,13.6) | 0.04 | <.001 |
| Rubidium | 8.45 (6.8,11.9) | 5.55 (4.16,6.76) | 9.32 (5.76,12.5) | 4.68 (2.98,6.20) | 0.002 | <.001 |
| Molybdenum | 0.75 (0.46,0.88) | <0.64 (LOQ) | <0.64 (LOQ) | <0.64 (LOQ) | - | - |
| Chromium | <0.48 (LOQ) | <0.48 (LOQ) | <0.48 (LOQ) | 0.60 (0.43,1.28) | - | - |
| Vanadium | <0.16 (LOQ) | <0.16 (LOQ) | <0.16 (LOQ) | 0.19 (0.14,0.47) | - | - |
| Cobalt | 0.08 (0.05,0.11) | 0.08 (0.06,0.13) | 0.10 (0.06,0.14) | 0.08 (0.07,0.14) | 0.007 | 0.28 |
| **TOTAL TRACE** | 427 (339,610) | 501 (365,634) | 478 (433,617) | 537 (438,820) | 0.60 | 0.13 |

### Compliance of foods with European legislation

Analysis of all feline or canine foods revealed broad non-compliance with all EU (FEDIAF) Guidelines; 94% (91/97) and 61% (46/80) of wet and dry foods, respectively failed to comply with all guidelines (Figure 2a,b). While a majority of foods complied with ≥8 of 11 guidelines (Dry food, 99% (79/80); Wet food, 83% (81/97)) many, particularly wet food, failed to either provide the minimum (e.g. for copper, 20 % of wet food; Figure 1c) or exceeded the maximum (e.g. for selenium, 76% of wet food; Figure 1d) levels of at least one essential mineral. Many had marked mineral imbalance (e.g. 29% of wet, 20% of dry for Ca:P ratio; Figure 1a) and some individual products had markedly variable mineral composition (e.g. five diets represented in Figure 2c). Repeated analysis of a subset of wet foods with a different batch-ID again showed broad noncompliance in line with the original dataset, indicating that our analysis was not an isolated single-batch effect (Figure 2d). Pictorial representation of compliance of all wet and dry foods analysed, according to FEDIAF guidelines is presented in Supplementary information (Figure S3, dry foods; Figure S4, wet foods).

**Figure 2:**
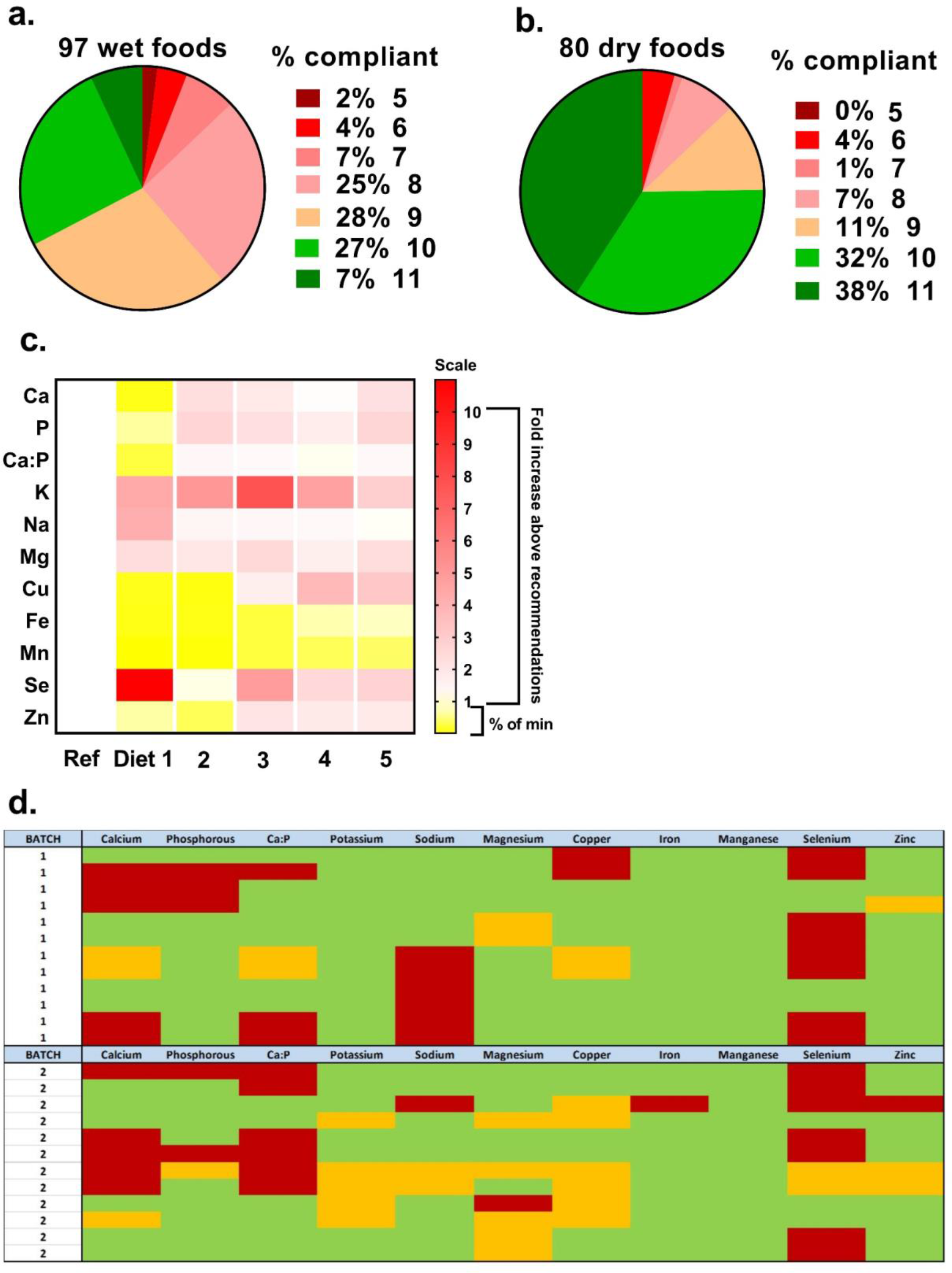
Compliance of complete, wet and dry pet foods with European guidelines. All wet (**a**) and dry (**b**) foods analysed were assessed against current European (FEDIAF) guidelines for individual elements (nutritional minimum and maximum). The percent of total that complied with all 11 of 13 guidelines (chloride and iodine were not assessed) are represented as green (e.g. 6 of 97 = 7% wet foods). **c**, Five individual diets are represented to show broad non-compliance and marked variability in elemental content. Ref, optimal nutritional content for each element (i.e. mid-way between nutritional minimum and maximum). Scale to 1 is percent below minimum (e.g. 0.20 is 5-fold less than nutritional minimum), scale 1-10 is fold-content above nutritional maximum. **d**, A subset of the original dataset (Batch 1) was purchased for a second time (Batch 2) with a different batch-ID to assess repeat compliance.

### Undesirable metal elements in food

Most undesirable metal elements in foods analysed (e.g. beryllium, thallium, lead, cadmium, uranium) were below detectable or quantifiable levels. However, some foods, particularly those with fish, were relatively high in the metalloid, arsenic (Figure 3a). Further examination revealed that only those foods that declared incorporation of higher levels of fish derivatives, as oppose to fish oil, had high levels of arsenic with three being above a safe upper limit (SUL) for organic As, two above the legal limit for food stuffs (Figure 3b). Indeed, As levels tended to increase steeply when ≥ 30-40% fish was declared (Figure 3c). Furthermore, a subset (× 10) of foods low or high in As were sent for direct analysis of mercury (Hg), which correlated well with As content (Figure 3d).

**Figure 3:**
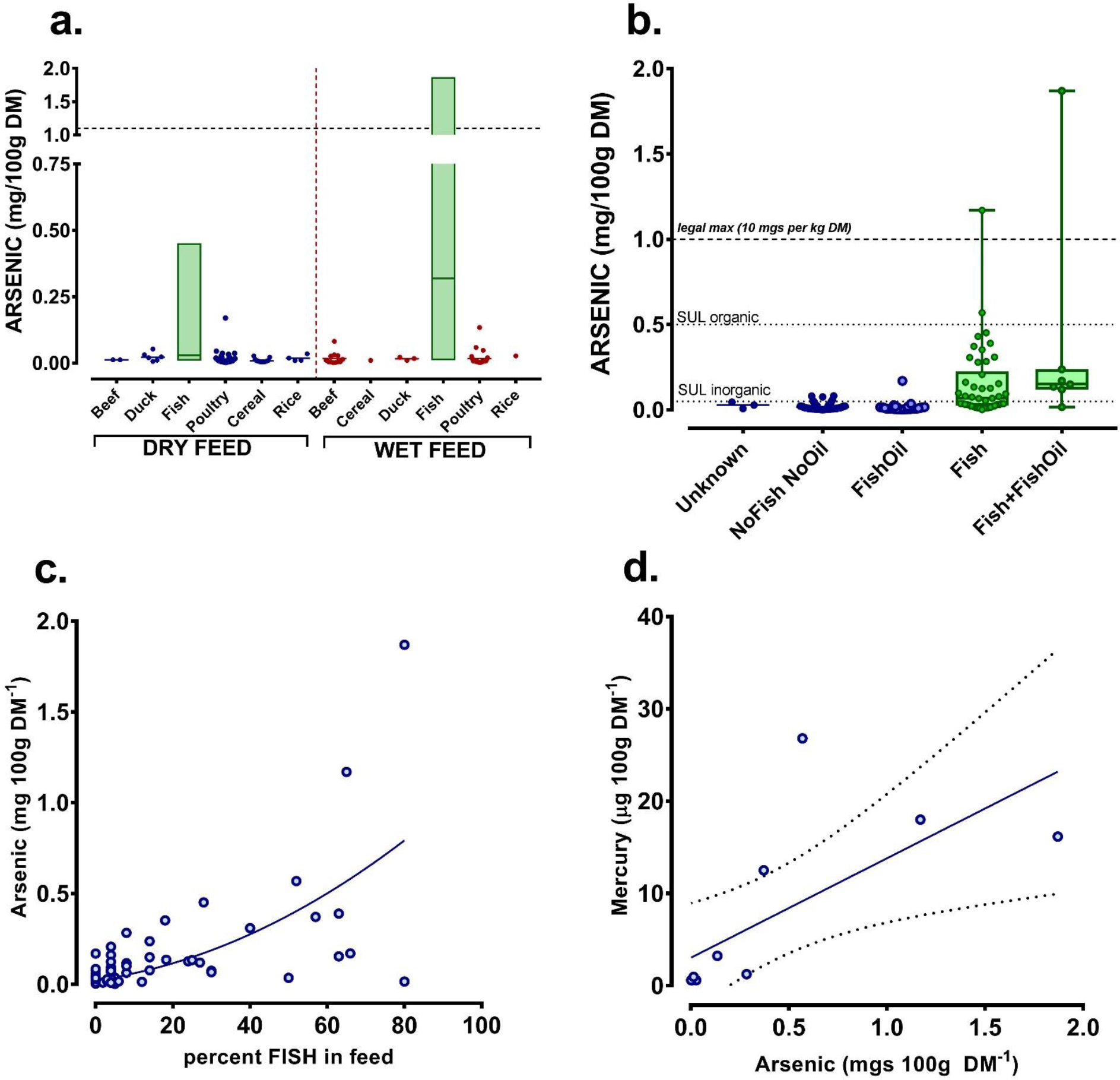
Arsenic in wet and dry pet food. Arsenic in wet and dry foods was analysed by ICP-MS (see Methods). **a)** Foods were classified according to main declared ingredient and are presented as dot-plots for visual clarity or box-plots (line at median, box represents range). Upper horizontal dotted line is legal maximum (10 mg/kg DM). Vertical dotted line separates dry from wet food. **b)** comparison of all foods declaring incorporation of fish derivatives (FISH) or fish oil (FishOil), none (NoFishNoOil) or both (Fish+FishOil). Three diets remained unknown as to fish or fishoil incorporation. **c)** scatterplot of the percent incorporation of fish derivatives vs. arsenic in food. Line is non-linear (quadratic) regression (r^2^, 0.47). **d)** Ten samples of food with low (×5) or high (×5) arsenic levels were sent for Direct Mercury Analysis (DMA-80) at The British Geological Survey (Keyworth, UK). Solid Line is linear regression with dotted lines 95% C.I. and the equation for the line is y=10.78^∗^X+3.031; F_1,9_=8.4, P=0.01).

## Discussion

This study, for the first time in the EU, has objectively and comprehensively assessed mineral composition and mineral balance of a large number of complete, wet and dry pet foods bought in pet-food supermarkets in the UK. The results suggest broad non-compliance with current EU guidelines for pet food, in that 92% of wet and 61% of dry foods did not comply with all recommendations. For a few individual products, mineral content often either far exceeded or did not meet nutritional requirements. A number had such unbalanced mineral content (e.g. Ca:P) that exclusive, longer-term feeding of these products could compromise general well-being of companion animals. Finally, we found that diets with high fish content also had relatively high arsenic content. Given that chronic arsenic intake in human epidemiological studies is associated with albuminuria, proteinuria and increased mortality from kidney disease (Zheng et al., 2014) then one must question why a high proportion of these diets are fed to domestic cats, whom are prone to chronic kidney disease (Lawson et al., 2015).

### Macronutrient and energy composition of the diets

On an as-fed basis (cats and dogs) the wet foods in our dataset had greater crude protein, fat, ash and moisture content than dry foods, with the latter having higher energy, fibre and carbohydrate (reported on the label as nitrogen-free extract [NFE]). Corrected for moisture content, energy density of wet and dry foods was similar. In accord with being obligate carnivores, feline diets had more crude protein that canine diets. A recent study estimated that feral felids consumed approximately 52% crude protein, 46% fat, 11% ash and only 2% N-free extract (Plantinga et al., 2011). These values, with the exception of fat, approximate to feline wet foods in this study (see Table 1) and are consistent with animals that would be engaging in far greater physical activity than a relatively sedentary housecat. Dry feline and canine pet food has a more carbohydrate-rich content than wet food. Canines have evolved away from their wolf ancestors, partially through adaptation toward consumption of a carbohydrate-rich, omnivorous diet (Axelsson et al., 2013). Felines, on the other hand, have remained remarkable true to their lineage as obligate carnivores from arid environments (Driscoll et al., 2007); the majority of their essential nutrients (e.g. arginine, taurine, arachidonic acid, retinol and niacin) can be obtained readily from eating other animals together with efficient gastrointestinal extraction and retention of water from solid food (Montague et al., 2014). Guideline feed amounts on pet food labels are on an as-fed (i.e. weight of food) basis. Owners can become confused by labelling information which may contribute to companion animal obesity, since dry foods that are predominantly fed have much higher energy density than wet foods (German, 2006). In addition, 40% (71 of 177) of foods analysed in our study had ≥10% ash on a dry matter basis, nine with ash ≥14%. Higher ash intake is associated with CKD in cats (Hughes et al., 2002).

### Compliance with EU guidelines

In 2010 a European scientific advisory board was established to ensure that current state-of-the-art knowledge on complete nutrition for health, well-being and longevity could be implemented for companion animals across EU states, via the industry’s relevant regulatory body, FEDIAF (FEDIAF, 2013). Their recommendations cover what is considered optimal intake of macronutrients such as protein and fat, micronutrients such as specific amino acids, trace elements and other minerals and vitamins to meet requirements for maintenance of good health. Minimum requirements are based on published values (NRC, 2006) or unpublished industry research. Nutritional maximums reflect the level in food that is not associated with any adverse effect, where known (FEDIAF, 2013). Consideration of these recommendations in light of our study where 11 of 13 recommendations were referenced, we found that the majority were broadly compliant (i.e. with ≥8 of 11 guidelines), particularly dry food, but many individual foods were far outside the recommended range (e.g. Ca:P ratio of 0.06 or 2.96:1). If fed over long periods of time, these products could compromise optimal health of companion animals (Schoenmakers et al., 2000).

Seventy-six percent of wet foods had greater than the maximum concentration (legal) of selenium, with seven having more than 3-fold the nutritional maximum (i.e. 3-fold more than 56.8 μg-100g DM^-1^). Selenosis is rare in companion animals (Burk and Hill, 2015), but nevertheless has been reported (Yu et al., 2006). We acknowledge that the current study has not assessed bioavailability of ingested selenium, which can be variable between companion animals, but rather has simply assessed concentration in the foods – for which the EU guidelines are applicable. Our highly variable results for total measured selenium in pet food do call into question the evidence base for designating such a tight recommended range. Some diets also had ≥2-fold the legal maximum for dietary copper (4.66 vs. recommendation, 2.80 mg-100g DM^-1^] or less than half the nutritional minimum (0.18 vs. recommendation, 0.50 mg-100g DM^-1^]. Similarly, although copper toxicity is also rare in companion animals (Wills and Simpson, 1994), excess copper in the diet can reduce bioavailability of zinc and iron (Gulec and Collins, 2014). Copper deficiency can directly impact reproductive performance and cause coat/hair instability (Zentek and Meyer, 1991). Relatively high levels of certain trace minerals can either lead to toxicosis *per se* or significantly affect bioavailability of other trace minerals through gastrointestinal interactions, in particular between calcium, iron, copper, zinc, magnesium and phosphate (Forbes, 1983). The mineral balance of complete diets is therefore important if they are to be fed exclusively over long periods of time. Excess calcium intake (≈ ≥2.5g·100g DM^-1^; 32 foods in our study) can lead to joint, limb and skeletal deformities (Hazewinkel, 1989).

### Arsenic

Whilst the primary outcome of this study was to assess mineral composition of a range of wet and dry pet foods with reference to compliance against EU recommendations, it did not escape our attention that certain other undesirable elements such as arsenic were particularly high in certain foods (see Figure 4), particularly in diets for domestic cats. Indeed, three foods were above World Health Organization recommendations for safe upper limits for organic arsenic, two were above the current legal maximum for animal feedstuffs. We estimate that, if fed exclusively, annual intake (assuming modified Atwater criteria for intake per kg metabolic bodyweight) for the five foods with highest arsenic content would yield a total exposure of between 800-6500 mgs arsenic per year. Arsenic is water soluble and accumulates in organs such as the liver and kidney, where it is relatively toxic either by itself (Yu et al., 2006) or via interactions with other catalytic metals such as iron and copper (Valko et al., 2005). High urinary arsenic is associated with increased risk of CKD (Chen et al., 2014). Certain soil and water sources are relatively contaminated with an abundance of arsenic (Robles-Osorio et al., 2015), due to its inclusion in agrochemicals such as phosphate fertilisers (Jayasumana et al., 2015). Arsenic may therefore bio-accumulate in the tissues of animals such as fish that inhabit these waters (Rosso et al., 2013) and in the animals that consume these fish sources such as the domestic cat. It remains to be determined if relatively high, long-term arsenic intake predisposes the domestic cat to chronic kidney disease or whether high intake of other metals that are also prevalent in fish may also have an effect, such as mercury (Storelli et al., 2007).

### Limitations of the study

Our analysis has not been exhaustive. Rather, we present a snap-shot of current popular foods bought in UK pet supermarkets. Although owners primarily feed their pets complete diets, a proportion of nutritional intake may come from complimentary foods – for which we have made no assessment of mineral composition. A number of surveys have indicated that up to 50% of dog owners and approximately 20% of cat owners supplement the daily ration with such treats (PDSA, 2013). However, complimentary foods are rarely nutritionally-balanced products, especially with respect to mineral composition. For example, high-protein ‘jerky’ supplements are virtually devoid of calcium and high in phosphorus.

By design, this study has not considered bioavailability of minerals. It is possible that a non-compliant diet with, for example, high copper or selenium does not cause copper or selenium toxicosis due to mineral or food interactions in the gastrointestinal tract that prevent absorption. Nevertheless, the EU guidelines, for which we assessed compliance against, refer to concentrations in foods taking into account margins of safety and thus should, at least, be adhered to. Mineral bioavailability and metabolism is complex. Gut absorption of microminerals such as zinc or copper can be markedly reduced if the diet has relatively high fibre or phytate content; high dietary calcium can inhibit gastrointestinal iron and zinc absorption (Forbes, 1983). For animals with sub-clinical disease, such as early-stage kidney disease, variation in hormonal status (e.g. parathyroid hormone) or vitamin status (e.g. Vitamin D) can profoundly influence mineral uptake from the gut (Llach and Massry, 1985).

In conclusion, this study is the first to sample a wide range of wet and dry pet foods, designated as ‘complete’ and widely available to consumers in the UK, for mineral composition. A majority were non-compliant according to current European recommendations (FEDIAF, 2013). Many had either insufficient, excessive or an inappropriate balance of minerals which, if fed exclusively for a long period of time, could underpin a host of clinical diseases in dogs and cats including skeletal, neurological, or dermatological disease. Furthermore, foods with relatively high levels of fish or fish derivatives (i.e. ≥ 14%) also had high levels of undesirable metal elements such as arsenic, which bioaccumulate in internal organs and may contribute toward a plethora of sub-clinical disease states (Ercal et al., 2001, Mertz, 1981). The data suggest a need for better compliance to current recommendations for mineral nutrition of companion animals in order to safeguard animal health, and for a better understanding of the mineral requirements of cats and dogs. For consumers, the study can offer a number of suggestions to limit the impact of such variability in pet foods: 1) weigh 100g of wet and/or dry food to grasp how much you are feeding and feed accordingly, 2) feed a diet of mainly dry with some wet food, 3) choose foods with a declared as-fed mineral content (i.e. % ash on the label) of not more than 10% for dry food (three of our foods exceeded this value) and not more than 2% for wet food (sixty of our foods exceeded this value), 4) make the feeding of foods with a high fish content a rarity rather than the norm and 5) vary the types of food fed over extended periods.

## Author contributions

D.S.G., C.W. and M.D. designed research; L.J., R.A., C.D., C.W. & D.S.G. conducted the research; D.S.G. and M.D. co-wrote the manuscript. All authors critically evaluated the paper. D.S.G. conducted the statistical analyses. D.S.G. and M.A.J.D. have primary responsibility for its final content.

